## Supplemental information for "Perceptual saccadic suppression starts in the retina"

Idrees†, Baumann†, et al.

### Perceptual saccadic suppression starts in the retina

Saad Idrees†, Matthias P. Baumann†, Felix Franke,
Thomas A. Münch∗, and Ziad M. Hafed∗

#### Supplementary information

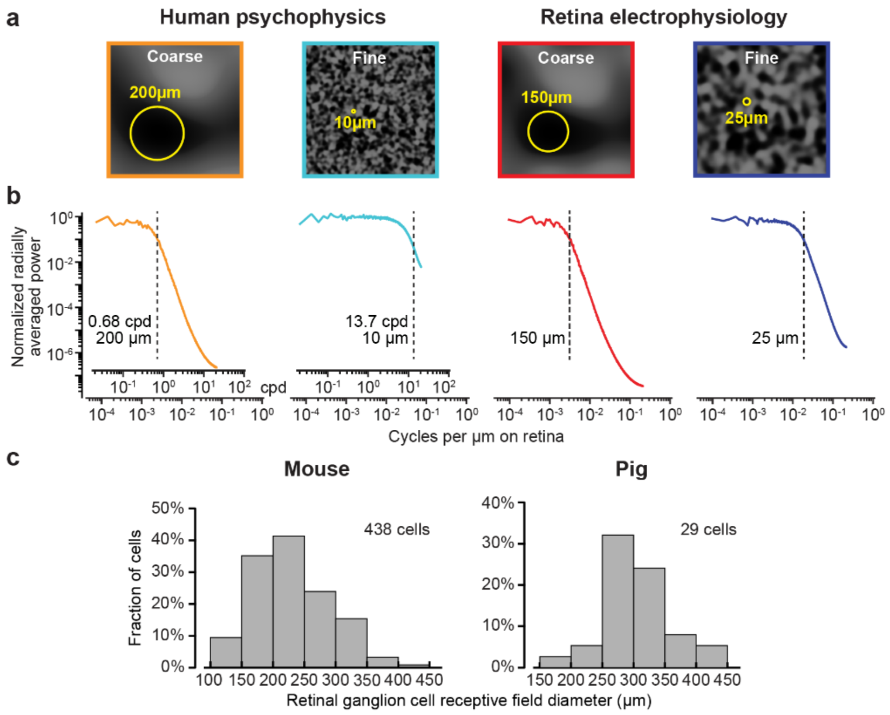


**Supplementary Figure 1 Textured backgrounds tailored to receptive field sizes of retinal ganglion cells (coarse) or bipolar cells (fine) in the different species that we studied. (a)** We created textures by convolving random binary pixel images with a Gaussian blurring filter. We varied the $\sigma$ parameter of the Gaussian blurring filter (Methods) to define a so-called spatial scale for the resulting texture (indicated as yellow circles in the examples shown). For each species, we picked the spatial scale to result in dark or bright image blobs that approximated the sizes of either retinal ganglion cell (coarse) or bipolar cell (fine) receptive fields, and we then set $\sigma$ to half the spatial scale value (Methods). **(b)** Radially-averaged power spectra for textures like in **a**, normalized to the maximum average power. Low-pass characteristics in all spatial scales were clear, as expected: less than 5% of the total average power was above the spatial frequency corresponding to the specific spatial scale of a given texture (vertical dashed lines). The inset x-axes in the first two spectra (used for human perceptual experiments) show units of cycles per degree (cpd) in addition to cycles per µm on the retina. **(c)** Histograms showing the distributions of receptive field diameters (Methods) in mouse (left) and pig (right) for a subset of retinal ganglion cells that we recorded. Since the distributions were generally similar, we used the same spatial scale parameter for the retinal recordings in both species. Human spatial scale parameters were estimated based on human receptive field diameters from the literature (Methods).

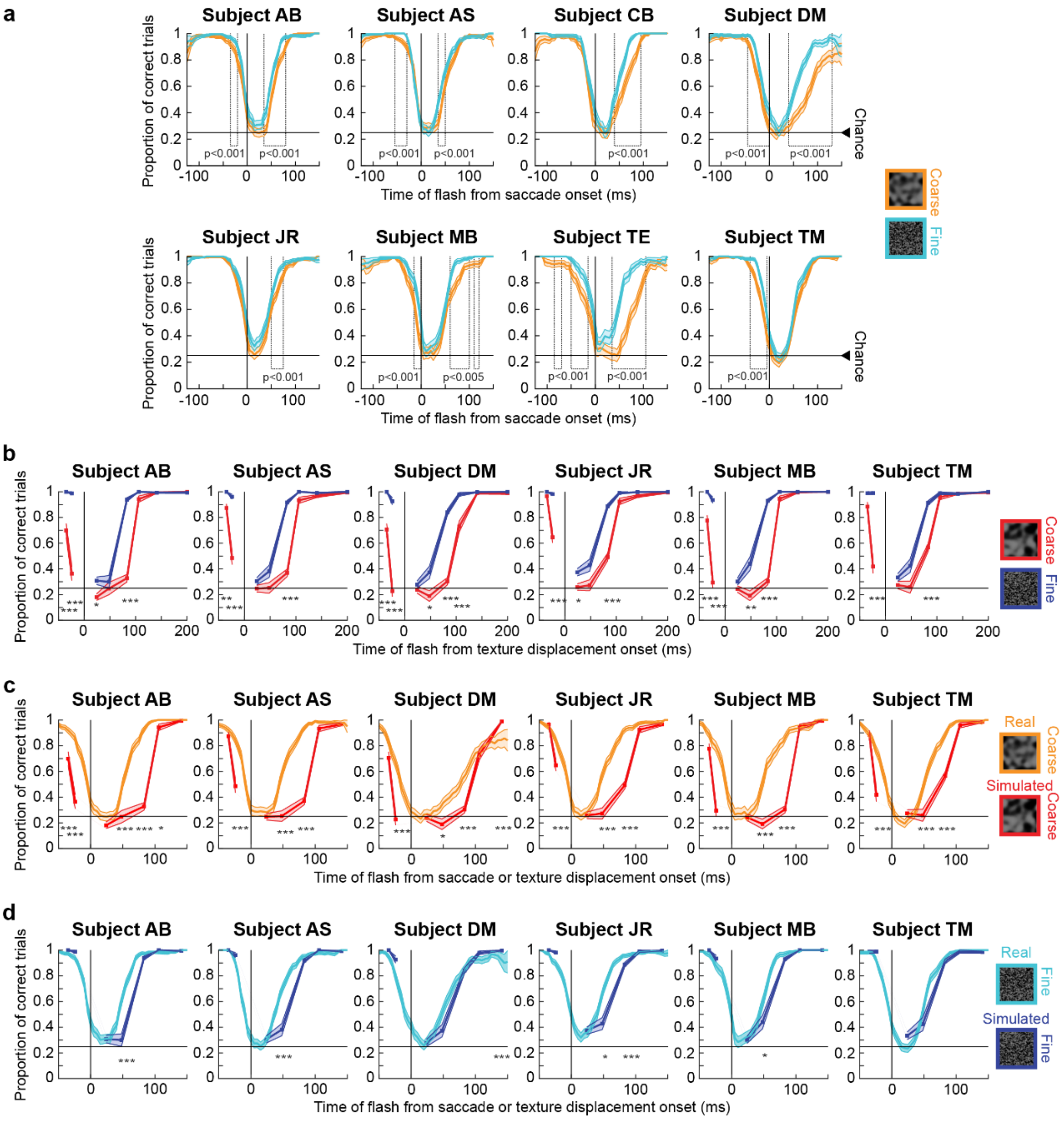


**Supplementary Figure 2 Individual subject results from the perceptual experiments of Figs. 1, 6. (a)** Identical analyses to Fig. 1d, shown separately for each individual subject. Error bars: s.e.m. across trials. All subjects experienced strong perceptual saccadic suppression, going from near-perfect localization performance to near-chance performance at peak suppression. Moreover, using strict statistical criteria (indicated in the figure and described in detail in Methods), all subjects had significant time clusters during which perception was different between saccadic suppression for saccades generated across coarse or fine textures. Also see Fig. 2 and Supplementary Fig. 4. **(b)** Same analyses as in Fig. 6d, but now showing individual subject results when saccades were replaced by saccade-like texture displacements during fixation. All subjects showed longer suppression after coarse texture displacements than after fine texture displacements; all subjects also showed earlier and stronger "pre-saccadic" suppression for coarse textures. Note that this "pre-saccadic" effect is purely visual, since the subjects never made saccades in this condition. Also, note that all subjects who participated in this experiment had also participated in the version with real saccades in **a**. Therefore, whether with or without saccades, perceptual suppression depended on image statistics. Also see Fig. 7 and

Supplementary Fig. 7. **(c, d)** Comparisons of perceptual suppression between real and simulated saccades across coarse **(c)** and fine **(d)** textures, as in Fig. 6e, f but now separating data from individual subjects. Note how even pre-saccadic suppression was prolonged in simulated relative to real saccades (i.e. started earlier in simulated saccades) in the coarse texture condition, which was most effective in causing suppression overall. Error bars: s.e.m. across trials. Asterisks in **b** denote a significant difference between coarse and fine conditions at the indicated flash time ($\chi^2$ tests with Bonferroni corrections; $*$ $p<0.005$, $**$ $p<0.001$, $***$ $p<0.0001$). Asterisks in **c**, **d** denote significant differences ($*$ $p<0.007$, $**$ $p<0.0014$, $***$ $p<0.00014$) between real and simulated saccades, comparing perception of a flash at the indicated time delay after simulated saccades to the corresponding time bin (+/- 25 ms) from the real saccade condition.

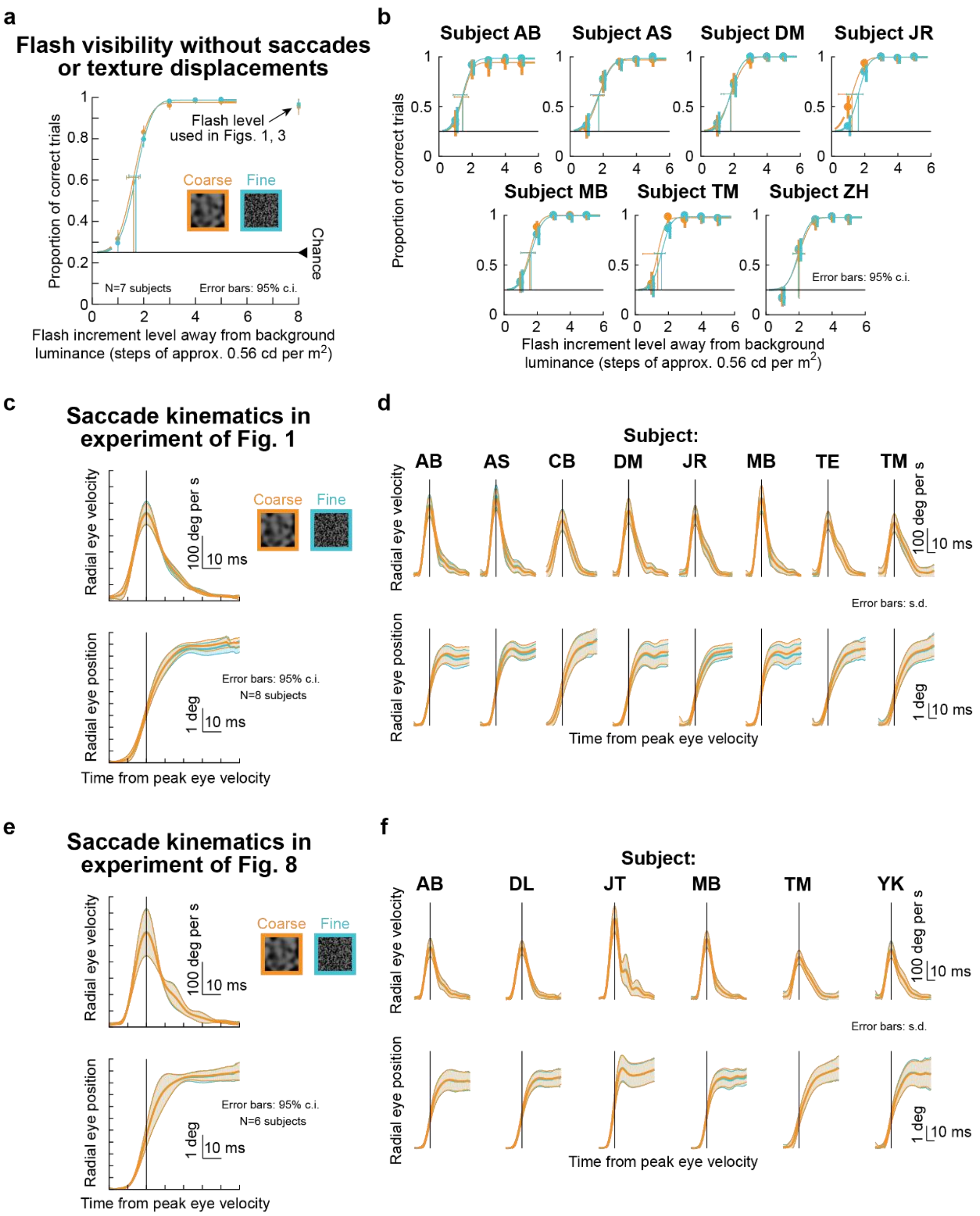


**Supplementary Figure 3 Controls for flash visibility and motor variability in our perceptual experiments of Figs. 1, 6, 8. (a)** For the same textures as in Figs. 1, 6, we asked subjects to maintain fixation. At a random time, a luminance pedestal appeared as in the main experiments (Figs. 1, 6), but this time, we varied its contrast across trials (Methods). We ensured that no microsaccades occurred near the flash onset time (Methods). Psychometric curves of localization performance indicate that, at the flash contrast used in Figs. 1, 6 (highlighted by the black arrow), subjects could easily detect flashes during simple fixation. Importantly, flash visibility was identical for coarse or fine textures at all contrasts. Therefore, flash visibility alone (or lack thereof) did not explain the main experiments' results (Figs. 1, 6). The strong perceptual suppression observed in Figs. 1, 6 was instead likely a function of interaction between visual transients associated with saccades or texture displacements and the flashes.

Also see Figs. 2, 7. **(b)** This idea is further supported by the fact that all individual subjects showed consistent results. All of these subjects had also participated in the experiments of Figs. 1, 6 (with the exception of subject ZH who only performed the control experiment). Psychometric curves were fit using the *psignifit 4 toolbox*[1], and error bars denote 95% confidence intervals. **(c)** We also checked for potential effects of motor variability on perceptual performance, in order to rule out the possibility that differences in performance between textures (Fig. 1) were due to differences in eye movement kinematics. For the experiments of Fig. 1, we plotted average radial eye velocity (top) and average radial eye position (bottom) across subjects (error bars denote 95% confidence intervals across the individual subjects' curves). There was no effect of background texture on movement kinematics. **(d)** This was also true for each subject. In this case, error bars denote s.d. across trials; saccade kinematics were not different when saccades were made across coarse or fine textures. **(e, f)** Same kinematic analyses, but now for the saccades of the experiment of Fig. 8. Scale bars are defined in their respective panels.

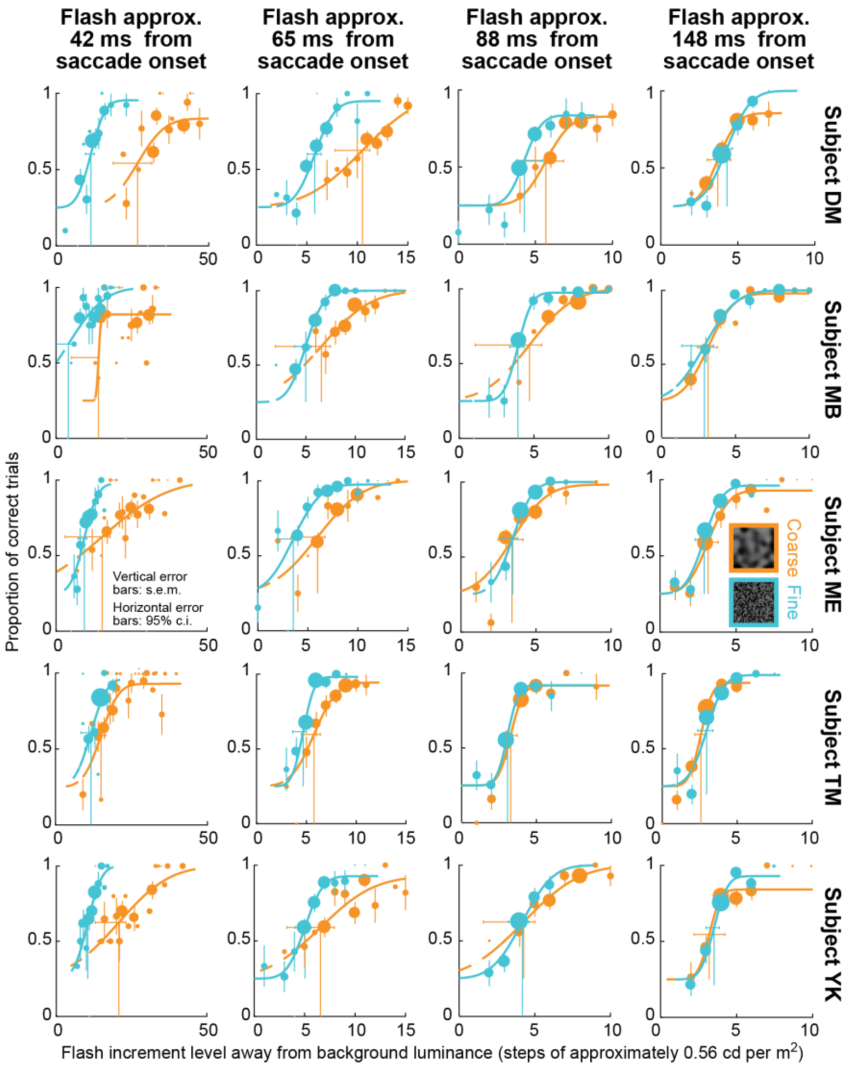


**Supplementary Figure 4 Individual subject results from the perceptual experiment of Fig. 2.** Each row shows psychometric curves like those shown in Fig. 2a-d, but for a single individual subject. Different rows show results from different subjects. The same conventions as in Fig. 2a-d apply. Here, we also scaled the size of each data point shown by the number of repetitions collected during the experiment. Note that we only show vertical error bars for data points with >10 repetitions, for clarity. Vertical error bars denote s.e.m. across repetitions of a given condition; horizontal error bars indicate 95% confidence intervals for the detection threshold of a given psychometric curve (i.e. the flash contrast resulting in threshold perceptual performance; Methods). Note that the x-axis ranges for the different columns (i.e. different flash times from saccade onset) are different from each other because of the varying amounts of perceptual saccadic suppression that occurred (Figs. 1-2). As can be seen, all subjects showed strong perceptual suppression near the time of saccade onset, with recovery occurring later in time, consistent with Fig. 1. Moreover, all subjects showed stronger perceptual suppression with coarse textures when compared to fine textures, again consistent with Fig. 1.

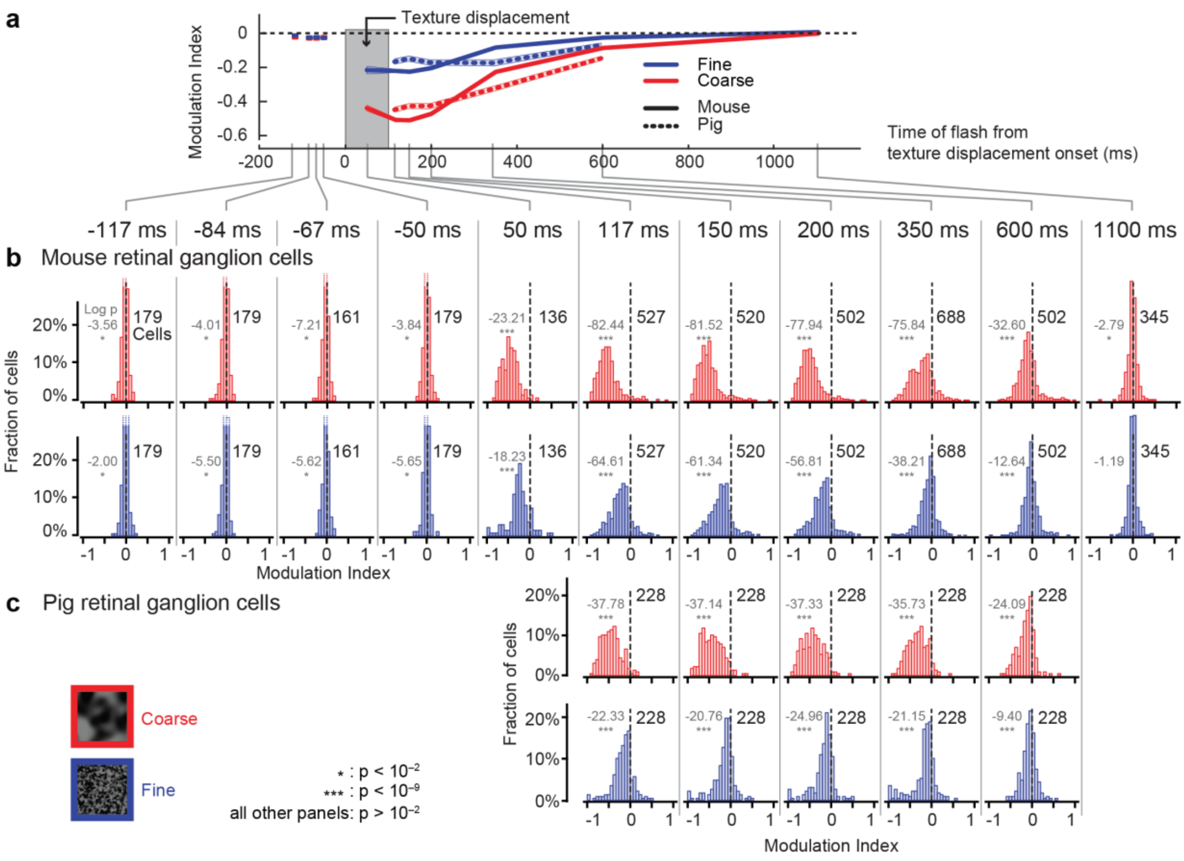


**Supplementary Figure 5 Population data detailing the properties of retinal "saccadic suppression". (a)** Replication of Fig. 3e, showing the time courses of retinal "saccadic suppression" in mouse and pig retinae. **(b, c)** Histograms of neuronal modulation indices for mouse **(b)** and pig **(c)** RGCs at different flash times relative to texture displacement onset. Red and blue denote coarse and fine textures, respectively. Black numbers in each panel indicate the numbers of RGCs analyzed for each condition; gray numbers show the logarithm (base 10) of the exact p-value (two-tailed Wilcoxon signed-rank test to determine if the population median was shifted away from 0). Asterisks additionally indicate the level of significance.

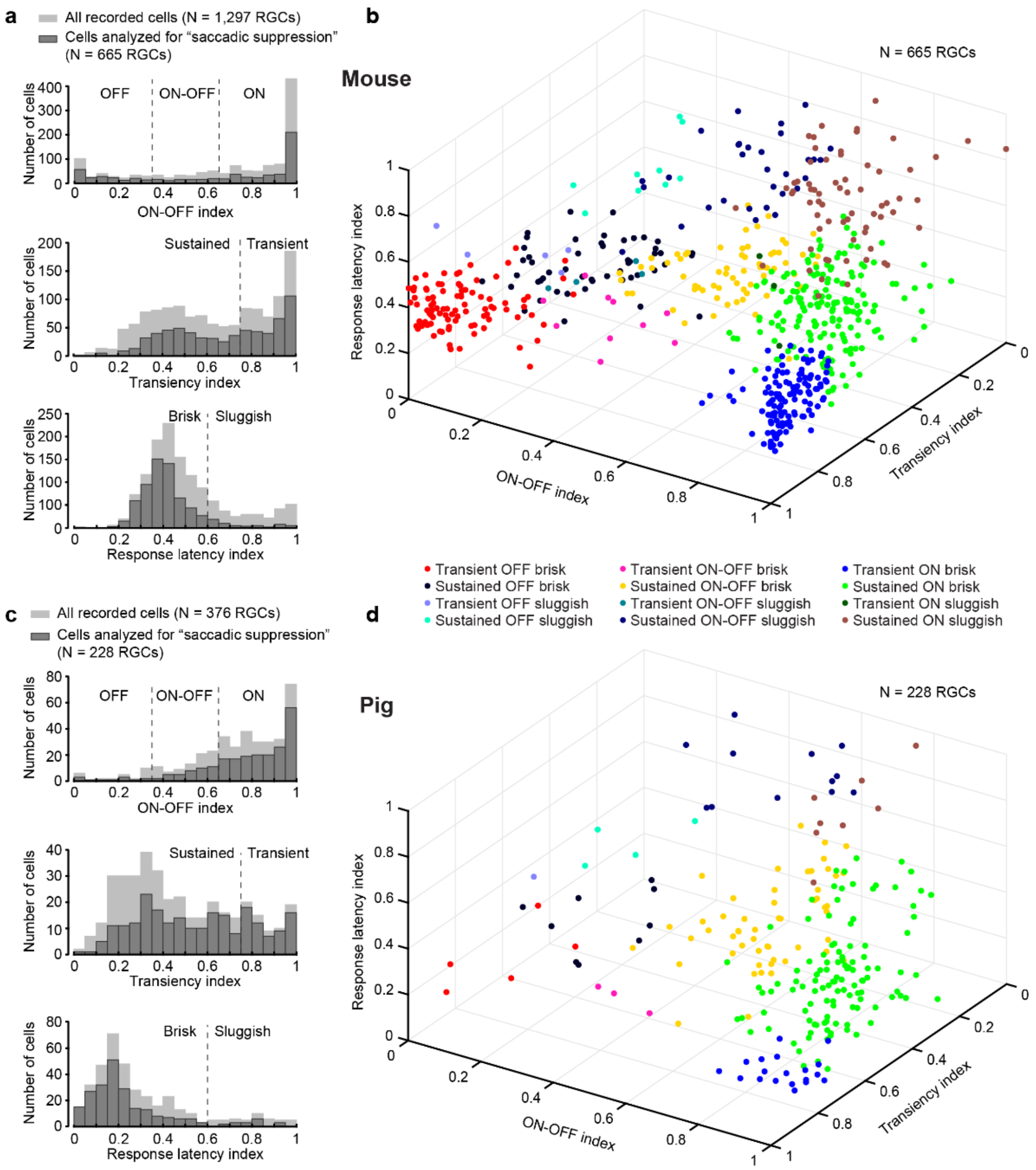


**Supplementary Figure 6 Diverse properties of RGCs included in our analysis.** We quantified the response properties of all recorded mouse **(a, b)** and pig **(c, d)** RGCs with respect to three neuronal response properties (see Methods): ON-OFF index, transiency index, and response latency index. Each histogram **(a**, **c)** was divided into 2 or 3 groups: RGCs could be OFF, ON-OFF, or ON (top histograms); transient or sustained (middle histograms); and brisk (short response latency) or sluggish (long response latency) (bottom histograms). Combined, this resulted in 12 response categories to which each recorded RGC belonged. The cells that could be analyzed for "saccadic suppression" and for which the response properties could be computed (dark gray histograms) spanned the entire range of response indices exhibited by all recorded cells for which these response properties were analyzed (light gray histograms). The three-dimensional scatter plots **(b**, **d)** show the projection of the RGC subsets considered in our analysis for "saccadic suppression" onto the 3 neuronal response indices. The 12 response categories, formed by the combination of histograms in **(a**, **c)**, can be seen in different colors.

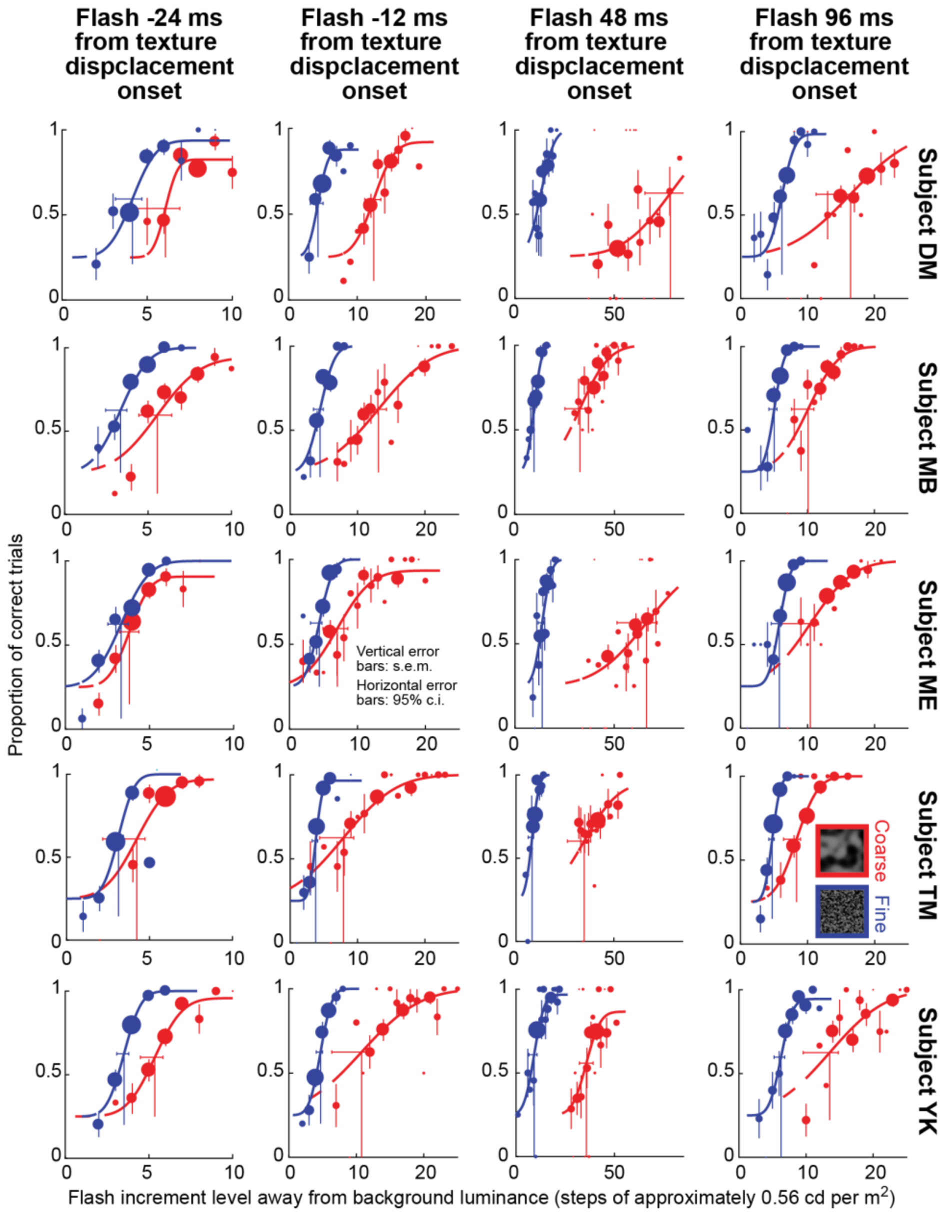


**Supplementary Figure 7 Individual subject results from the perceptual experiment of Fig. 7.** Same as Supplementary Fig. 4, but now for the experiment of Fig. 7. All subjects showed similar results: there was strong perceptual suppression before and after texture displacements in the absence of saccades, and the suppression effect was stronger when the displaced texture was coarse rather than fine. Note that in this experiment, we added an additional time sample prior to texture displacement onset, in comparison to Fig. 6, in order to demonstrate the robustness of this pre-displacement effect, and also to demonstrate the continuity of perceptual suppression in a time-locked fashion to texture displacement onset (Fig. 6).

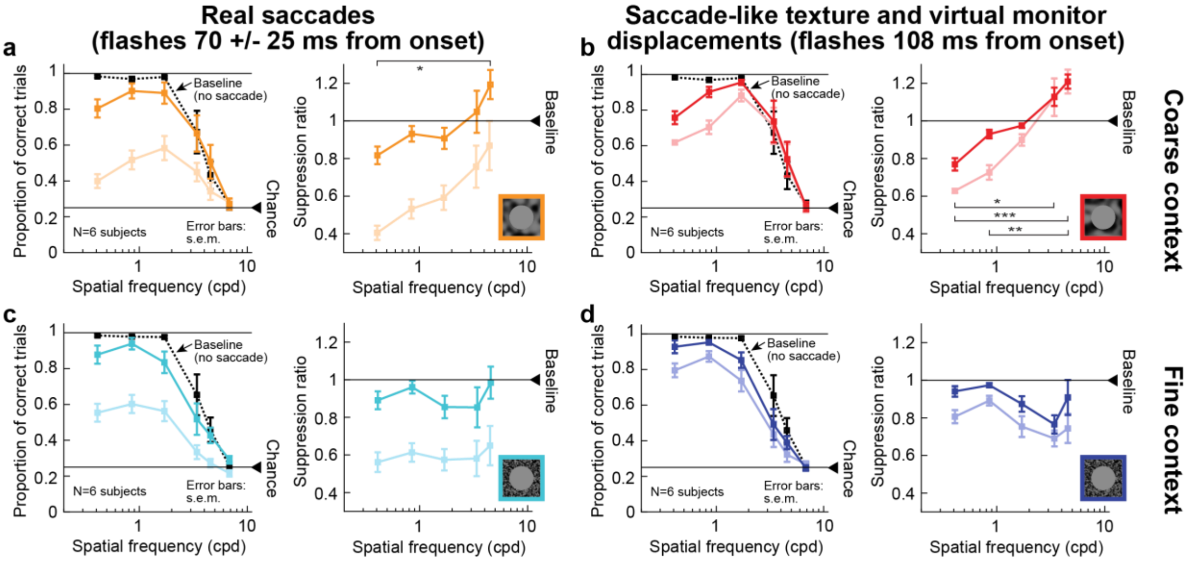


**Supplementary Figure 8 Recovery, with time, of perceptual suppression with real and simulated saccades in the experiment of Fig. 8. (a)** Same analysis as in Fig. 8b, but at a later time point of grating flash onsets relative to saccade onset. Faint curves show the data from Fig. 8b for comparison. At around 70 ms after saccade onset, perceptual recovery from saccadic suppression emerged, but the selectivity of suppression across different spatial frequencies was still present (there was a main effect of spatial frequency on suppression ratio; $\chi^2$=11.4, p=0.022, df=4, Kruskal-Wallis test). All other conventions are as in Fig. 8b ($*$ p<0.05, post-hoc pairwise test between the lowest and highest spatial frequencies). **(b)** Same analysis as in Fig. 8d, but at a later time point of flash onset after virtual monitor and texture displacement. The same observations as in **a** were made: perceptual recovery occurred at the later time point, but selectivity of suppression was still obvious ($\chi^2$=25.26, p<0.0001, df=4, Kruskal-Wallis test, same post hoc conventions as in Fig. 8d). The faint curves show the data from Fig. 8d for comparison. Note how this condition of displacements of the virtual monitor and texture surround resulted in longer lasting suppression than with real saccades (also see Supplementary Fig. 9). **(c, d)** Same analyses as in **a** and **b**, but with a fine texture surrounding the virtual monitor (Fig. 8e, f). Error bars in all panels denote s.e.m. All other conventions are as in Fig. 8.

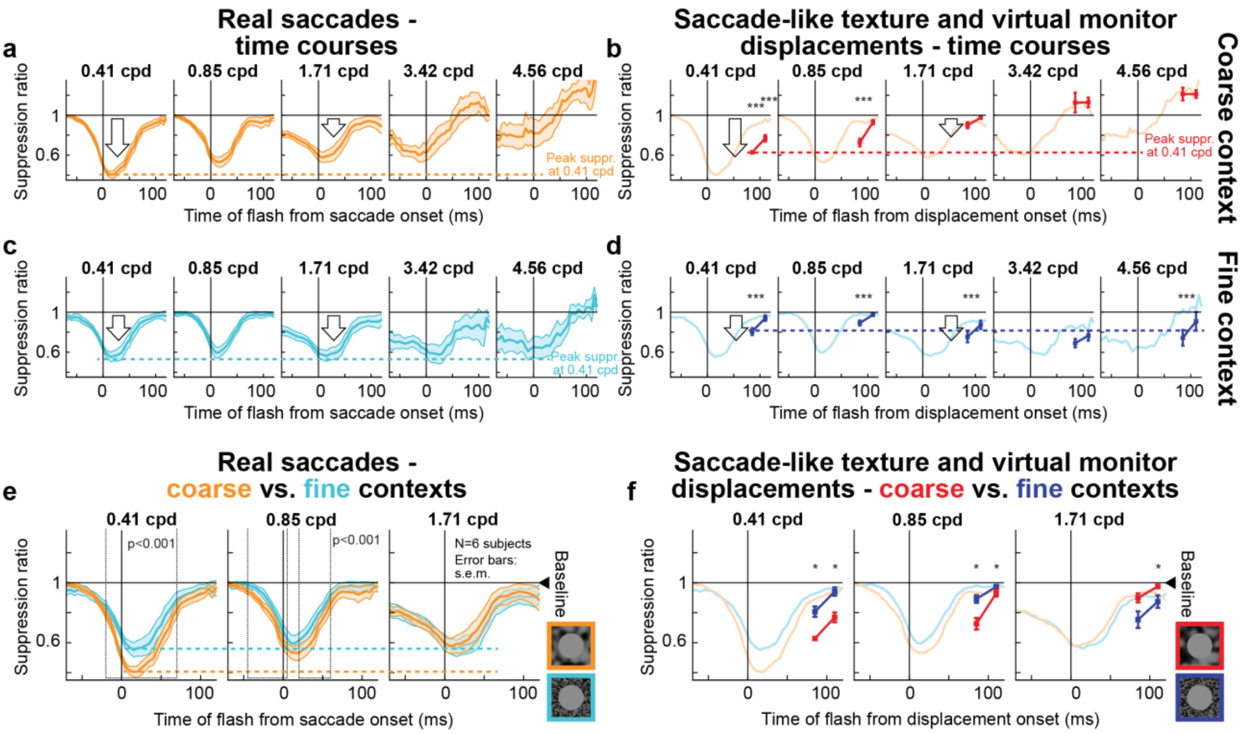


**Supplementary Fig. 9 Time courses of perceptual suppression with real and simulated saccades, as well as coarse and fine textures, in the experiment of Fig. 8. (a)** Time courses of suppression from Fig. 8a, b with a coarse surround around the virtual monitor. We used similar binning procedures to Fig. 1. Peak suppression was strongest when 0.41 cpd gratings were flashed and progressively weakened for higher spatial frequency gratings (horizontal colored dashed line across panels). **(b)** With simulated saccade-like virtual monitor and texture displacements, we sampled two grating flash times relative to displacement onset. Recovery at the later time point for each grating spatial frequency was evident. Moreover, selectivity of suppression as a function of grating spatial frequency was evident (horizontal colored dashed line across panels demonstrating the peak suppression for the lowest spatial frequency). The faint curves show time courses from **a** for comparison. Note how simulated saccades caused longer-lasting suppression than real saccades, exactly as in the experiment of Fig. 6 (∗∗∗ $p<0.0001$, $\chi^2$ tests with Bonferroni corrections comparing perceptual suppression with the simulated condition to a corresponding time bin in the real condition). **(c, d)** Similar analyses for fine texture surrounds around the virtual monitor. In this case, suppression was the same across all spatial frequencies (horizontal colored dashed lines across panels). **(e)** For real saccades, and for low spatial frequencies of gratings (i.e. when both coarse and fine surround contexts were associated with strong saccadic suppression), the coarse surround was associated with longer lasting suppression than the fine surround. This is consistent with the results of Fig. 1 when saccades were generated across full-screen textures. **(f)** This texture-dependence was also true with simulated saccades (∗ $p<0.05$, random permutation test comparing coarse and fine textures at a given grating flash time). Error bars in all panels denote s.e.m. All other conventions are as in Figs. 1, 6, 8.

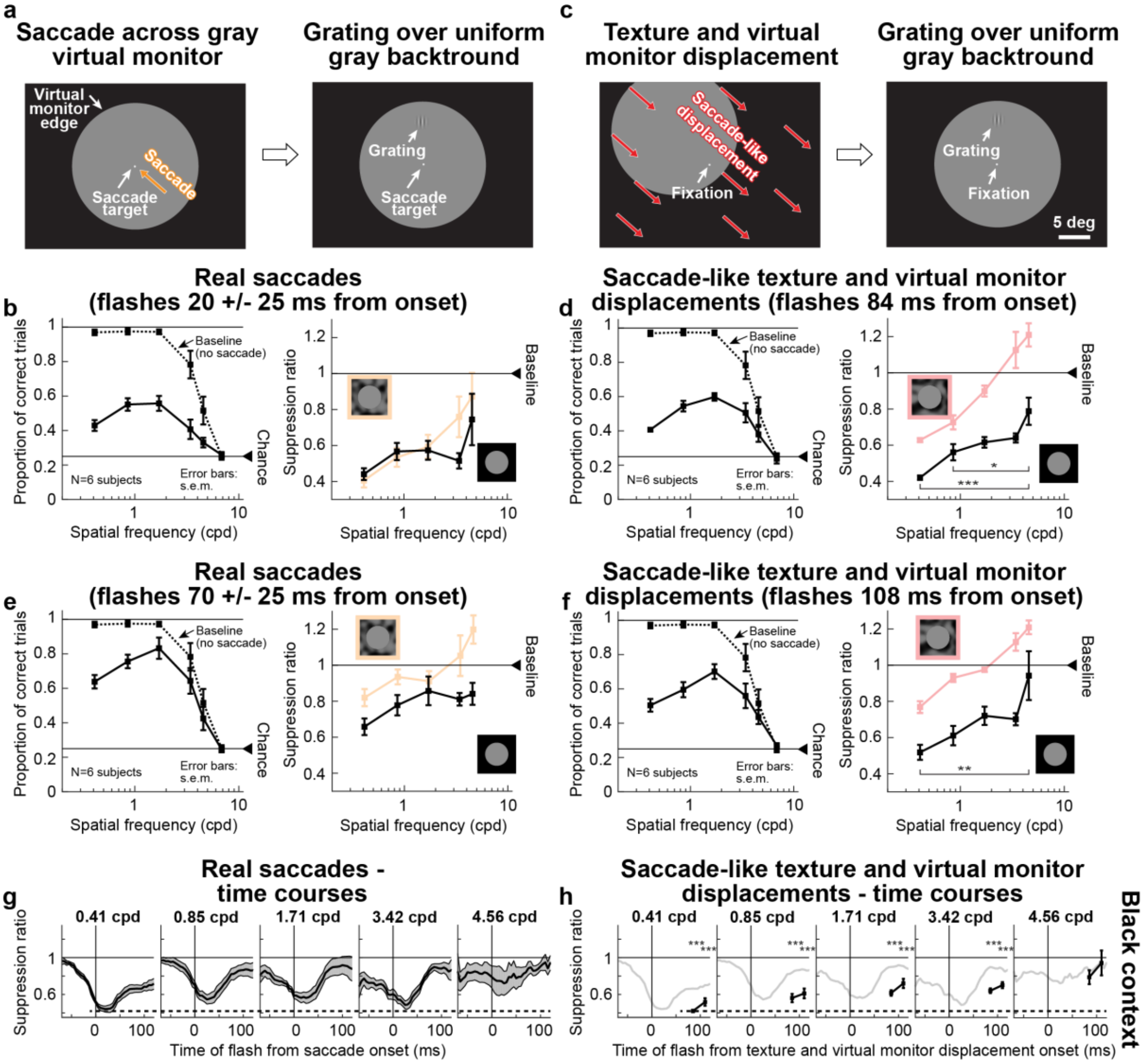


**Supplementary Fig. 10 Replicating the results of Fig. 8 but with black surrounds around a uniform gray display. (a)** We repeated the same experiment as in Fig. 8a but this time using a black surround around the virtual monitor, as we might normally do in experiments on saccadic suppression[2,3]. Note that a black (or white) surround is theoretically equivalent to an infinitely coarse surround; hence, we expected observations more similar to Fig. 8a-d (i.e. selectivity of suppression for low spatial frequencies) than Fig. 8e, f. **(b)** Similar suppression selectivity for low spatial frequencies occurred with real saccades as in Fig. 8b (faint curves replicate that data for comparison). **(c)** Same experiment as in Fig. 8c, but with a black surround. **(d)** Selectivity of suppression for low spatial frequencies was even more evident with simulated saccades. Faint curves show results from Fig. 8d for comparison. **(e, f)** Similar analyses at a later time point, identical to Supplementary Fig. 8. There was recovery for both real (**e**) and simulated (**f**) saccades (faint colored curves show data from Supplementary Fig. 8a, b at the same time points for easier comparison). Note that with black surrounds, suppression strength was larger overall than with either coarse or fine texture surrounds (as if the black surround was indeed an extension of the coarseness of the texture). **(g, h)** Full time courses of suppression as in Supplementary Fig. 9. All error bars denote s.e.m., and all conventions are similar to Fig. 8 and Supplementary Figs. 8, 9.

1 Schutt, H. H., Harmeling, S., Macke, J. H. & Wichmann, F. A. Painfree and accurate Bayesian estimation of psychometric functions for (potentially) overdispersed data. *Vision Res* **122**, 105-123 (2016).

2 Hafed, Z. M. & Krauzlis, R. J. Microsaccadic suppression of visual bursts in the primate superior colliculus. *J Neurosci* **30**, 9542-9547 (2010).

3 Chen, C. Y. & Hafed, Z. M. A neural locus for spatial-frequency specific saccadic suppression in visual-motor neurons of the primate superior colliculus. *J Neurophysiol* **117**, 1657-1673 (2017).
